# CNN-based Encoding and Decoding of Visual Object Recognition in Space and Time

**DOI:** 10.1101/118091

**Authors:** K. Seeliger, M. Fritsche, U. Güçlü, S. Schoenmakers, J.-M. Schoffelen, S. E. Bosch, M. A. J. van Gerven

## Abstract

Deep convolutional neural networks (CNNs) have been put forward as neurobiologically plausible models of the visual hierarchy. Using functional magnetic resonance imaging, CNN representations of visual stimuli have previously been shown to correspond to processing stages in the ventral and dorsal streams of the visual system. Whether this correspondence between models and brain signals also holds for activity acquired at high temporal resolution has been explored less exhaustively. Here, we addressed this question by combining CNN-based encoding models with magnetoencephalography (MEG). Human participants passively viewed 1000 images of objects while MEG signals were acquired. We modelled their high temporal resolution source-reconstructed cortical activity with CNNs, and observed a feedforward sweep across the visual hierarchy between 75-200 ms after stimulus onset. This spatiotemporal cascade was captured by the network layer representations, where the increasingly abstract stimulus representation in the hierarchical network model was reflected in different parts of the visual cortex, following the visual ventral stream. We further validated the accuracy of our encoding model by decoding stimulus identity in a left-out validation set of viewed objects, achieving state-of-the-art decoding accuracy.

## 1 Introduction

Automatic object recognition has been a long-standing and difficult research problem in computer vision. A few years ago, driven by the availability of large-scale computing and training data resources, automated object recognition has reached and surpassed human-level performance. The most successful object recognition models by far are deep feed-forward convolutional neural networks (CNNs), which can learn statistical properties of structured data such as natural images well (Schmidhuber, 2014).

CNNs can be seen as an abstraction of rate-based coding in biological neural circuits (Dayan and Abbott, 2005) and are inspired by the anatomical wiring of the visual system as a hierarchy of processing stages (Felleman and Van Essen, 1991). Receptive field properties become increasingly complex higher up in this visual hierarchy. That is, receptive fields in striate cortex respond to oriented bars in the visual input (Hubel and Wiesel, 1959), whereas receptive fields in inferior temporal cortex respond specifically to complex object properties. This is very similar to CNNs, which learn simple edge detectors in early layers and more abstract object features in higher layers. CNN-like architectures have been motivated from a neuroscientific point of view with the introduction of the Neocognitron by Fukushima (1980). This early model was already invariant to scale, translation and deformation; three major requirements of object recognition.

Another biological inspiration of contemporary CNNs is that their feature detectors (receptive fields) are learned from natural data (the natural environment) instead of using hand-engineered (a-priori) detectors. In addition, CNNs are capable of learning other types of feature detectors that are not considered by hand-designed (e.g. purely Gabor) models, but appeared in theoretical contemplations about visual system representations (e.g., see (Zeiler et al., 2010)). Similarly, in biological systems even the earliest cortical receptive fields will only exist if the organism has been exposed to certain visual environments in a critical learning period after birth (Blakemore and Cooper, 1970; Hubel and Wiesel, 1970). Using a universal learning algorithm throughout the hierarchy, almost all of these object recognition networks learn Gabor-like feature detectors in their lowest layer, similar to what is known about the response properties or neuronal populations in V1 (Hubel and Wiesel, 1959).

Comprehensive reviews of the use of state-of-the-art neural networks for probing neural information processing can be found in (Kriegeskorte, 2015; van Gerven, 2017; Yamins and DiCarlo, 2016). While some studies have also focused on the dorsal visual stream (Güçlü and van Gerven, 2015b; Eickenberg et al., 2016) and auditory stream (Güçlü et al., 2016) representations, most CNN-based investigations of neural processing have focused on understanding representations in the visual ventral stream (Yamins et al., 2014; Khaligh-Razavi and Kriegeskorte, 2014; Güçlü and van Gerven, 2015a). The similarities between biological and CNN object representations have mainly been studied with functional magnetic resonance imaging (fMRI), using either the encoding model framework or representational similarity analysis (RSA).

However, object recognition is a rapid process: discriminative information exists as early as about 100 ms after stimulus onset (Thorpe et al., 1996; van de Nieuwenhuijzen et al., 2013; Clarke, 2014). The seconds-long temporal delay and indirect nature of the fMRI BOLD signal therefore prohibit the investigation of how the network hierarchy sequentially represents stimulus features across time and throughout anatomical structures. An alternative approach is to combine CNNs with electrophysiological measurements of neuronal activity such as electroencephalography (EEG) or magnetoencephalography (MEG) to probe the dynamics of object processing in the human brain. This approach was pioneered by Cichy et al. (Cichy et al., 2016; Cichy and Teng, 2016), where RSA was used to show a correspondence between CNN layer representations and neural representations across space in fMRI and time in MEG.

In this study we expand on this work and used the encoding model framework to probe how CNN-based representations are expressed in space and time across the cortical surface using MEG. We encoded source-reconstructed brain activity in response to the single-pass presentation of a large, varied set of object images using the VGG-S CNN architecture (Chatfield et al., 2014), pretrained on the `imageNet` competition database. We show that the different layers of the CNN hierarchy can be mapped onto separate regions in the visual cortical hierarchy in space and time, and are able to model neural activity developed as early as 45-75 ms after stimulus onset. Furthermore, we demonstrate that the developed encoding model can be used to identify (decode) the perceived stimuli from MEG measurements, even at the single-trial level and in the absence of visual input. This indicates the possibility of future investigation of imagery-related processing (Horikawa et al., 2013; Horikawa and Kamitani, 2016; Naselaris et al., 2015).

## 2 Methods

Here, we first give a brief overview of the ideas behind our experimental design, followed by a description of data preprocessing, source modeling, anatomical alignment, the feature transformation of the stimuli and our encoding and decoding procedures in complete detail.

### 2.1 Experiment overview and rationale

In brief, we presented 1000 images of well-identifiable natural objects to each of 15 participants while we recorded the magnetoencephalogram. A random subset of 50 images was repeated 10 times (*validation set*), i.e. once in each block of 145 images, while the majority of 950 images was presented once (*estimation set*). There was no specific task, and the participants were asked to passively view the images. The design of each 3 seconds-long trial is illustrated in Figure 1. The extended post-stimulus fixation period allowed us to study sustained representations and decodability in the absence of the stimulus. The validation set was used to obtain a cleaner estimate of the event-related fields. Focusing on the variety of stimuli comes at the cost of not being able to obtain clean event-related fields for all of them, but is generally more beneficial for sampling as much of the feature space used for estimating the encoding model as possible given a relatively short experimental session.

**Figure 1:**
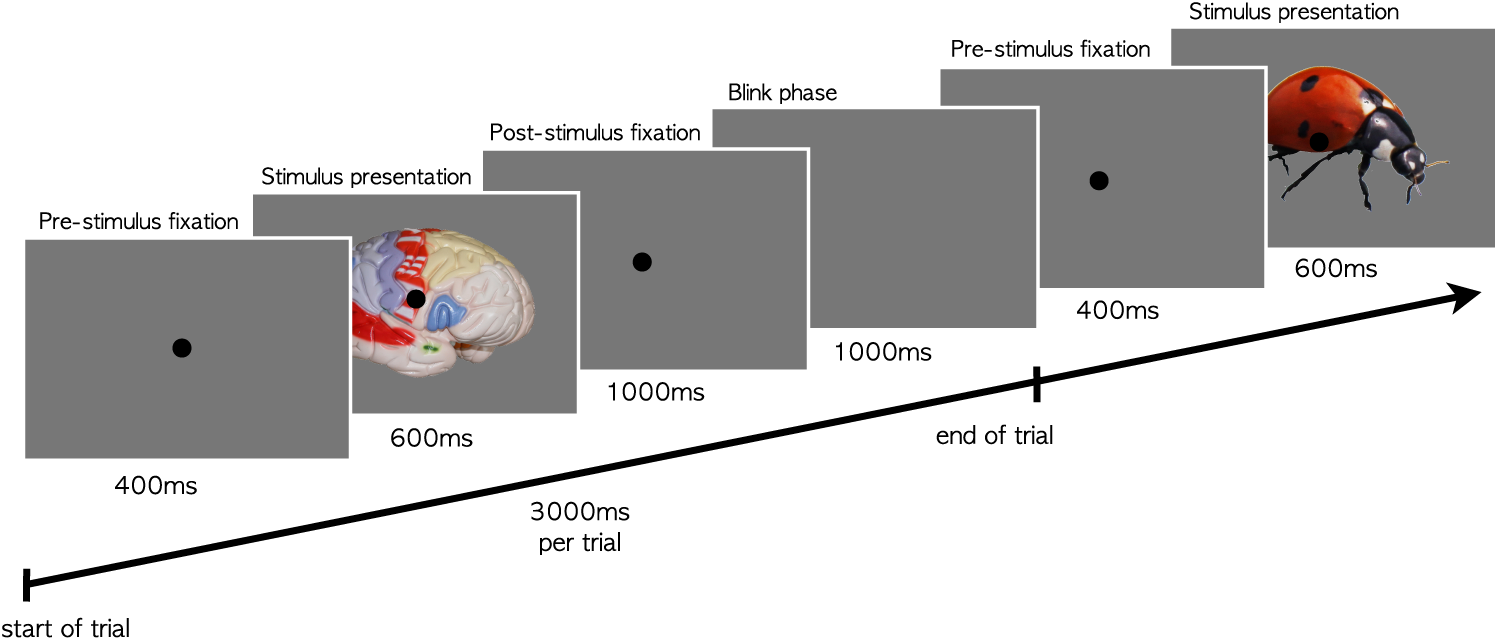
Design of individual trials. Images of objects were presented for 600 ms, preceded and followed by fixation periods. Participants could blink between each fixation. The experiment consisted of 1450 trials of 3 s length, resulting in 72 minutes of MEG acquisition time per participant.

The experimental design and data analysis follows the idea of system identification via encoding models (Wu et al., 2006; Gallant et al., 2011; Naselaris et al., 2011). Naturalistic stimuli were chosen under the assumption that sensory systems have evolved to be optimally adapted to the specific statistics of natural scenes (Bell and Sejnowski, 1997). The computational goal of arriving at invariant cortical representations of objects is believed to be realized by transforming the retinal input through a series of nonlinear processing steps in the ventral stream. The resulting representations are assumed to be reflected in characteristic patterns of brain activity and to be discoverable by using candidate representations to predict brain activity within an encoding model framework. Candidate representations were taken from the learned hierarchy in a state-of-the-art feedforward CNN (VGG-S from (Chatfield et al., 2014)), trained on natural photos with the objective function of classifying objects with perfect accuracy. Similar to what we know about biological visual systems, these networks start their feature detection hierarchy with low-level edge detectors and arrive at an invariant object representation. In-between these steps are intermediate learned representations of rising complexity.

To link neural networks to patterns of brain activity, we modelled cortical source activity from the MEG sensor activity using source reconstruction on individual brain surface models. Next, for each participant we developed a source-wise encoding model, predicting the modelled source activity globally using CNN representations. The encoding model uncovers when and where cortical activity is similar to these representations. The resulting encoding model can also be used for decoding, where the most likely stimulus is selected from a set of candidate stimuli based on observed brain activity.

### 2.2 Experimental design

#### 2.2.1 Stimuli

The presented natural images were mainly selected from the Bank of Standardized Stimuli (BOSS), and partly from Amsterdam Library of Object Images (ALOI) databases ( (Brodeur et al., 2010, 2014; Geusebroek et al., 2005) ) and from the author’s own collections. The BOSS database was standardized towards non-ambiguous identification or naming agreement. We chose a subset of images that could be classified correctly or similarly within the set of five most probable output classes of the VGG-S neural network. With the aim of obtaining 1000 classifiable stimulus images, we took 905 images from the BOSS database and chose the remaining 95 from the ALOI database and the author’s collections. For all images, the monochrome white or black background was changed to middle grey to reduce visual strain. From the 1000 images, a random subset of 950 images was presented once and will be referred to as the *estimation set.* Its function was to sample the representation spaces as comprehensively as possible in an experiment using a wide range of different natural stimuli. The remaining 50 images were repeated 10 times throughout the experiment and are referred to as the *validation set.* It amounted to approximately one third of the presented stimuli and was used to obtain event-related fields (ERFs) with a higher signal-to-noise ratio in order to report encoding and decoding performances. When using the validation set, we averaged single-trial source time courses over up to 10 trials of a stimulus, i.e. those that were not rejected during preprocessing. Similarly, the *first presentation* of an image of the validation set (a single trial from this set) was its first trial that was not rejected. The validation set was held out from training, and significance or the optimal layers were determined by cross-validating on the estimation set.

#### 2.2.2 Trial design and experiment regime

Each three-seconds long trial contained the presentation of one stimulus image for 600 ms, which was presented spanning 8 degrees of the visual field together with a black fixation point in the centre. Before and after the presentation there were phases of 400 ms (partly used as activity baseline) and 1000 ms (post-stimulus period) respectively in which just the fixation point was shown. Participants were allowed to blink within a 1000 ms period after each trial that was indicated by the absence of a fixation point. The total inter-stimulus interval was 2400 ms. See Figure 1 for the trial structure.

The experiment consisted of 10 blocks 145 images each, with a block duration of 435 s. Each block contained one complete repetition of the 50 images in the validation set. The validation set images were randomly permuted between the block’s estimation set images. The sequence was different for every participant. After each block we talked to the participant, allowing time for rest.

### 2.3 Experimental procedure

#### 2.3.1 Participants

We recruited our participants through the Research Participation System of Radboud University. The recruitment system prescreening excluded participants with any kind of permanent metallic implants or components, as well as those with brain-related health problems. We acquired recordings of 15 participants (3 male) aged 20 to 37 years (mean age: 25.2, median: 24).

The participants gave written informed consent in accordance with the Declaration of Helsinki. The study was approved by the local ethical review board (CMO region Arnhem-Nijmegen, The Netherlands) and was carried out in accordance with the approved guidelines.

#### 2.3.2 MEG system set-up

Participants were sitting in a 275-channel whole-head MEG system (CTF Systems Inc., Port Coquitlam, Canada) in a dark magnetically shielded room. We recorded the magnetoencephalogram at 1200 Hz. The stimuli were presented using an LCD projector, which lead to a quasi-instantaneous presentation with less than 1 ms delay. To maintain magnetic shielding, the projector was set up outside the magnetically shielded MEG scanner room, and the image was back-projected onto a translucent screen inside the scanner room with two mirrors. To present the experiment we used the stimulus software `Presentation` (Neurobehavioral Systems, 2016, https://www.neurobs.com) on a `Windows 7` system, with cautious system settings such as reduced network communications. MEG and additional data recordings (stimulus triggers, electro-oculogram, eye tracker data) were collected in multi-channel data files on a real-time `CentOS` system. Head localization was done on this system as well. Head tracking for manual position correction was done on an additional `NeuroDebian` system in `Matlab` with the algorithm described in (Stolk et al., 2013).

#### 2.3.3 Participant preparation

Before the experiment participants received information about the nature of the data recording modalities and reasons for avoiding movements and metallic objects in the scanner. They were asked to wear contact lenses in case of myopia, not to wear eye make-up on the MEG recording day and avoid scheduling MRI recordings during the previous few days. Participants were provided with non-magnetic clothes.

Electrodes were attached to the right collarbone and lower left rib, recording the electrocardiogram (ECG) to facilitate removing the heart signal noise in the MEG. Saccades and eye blinks were measured with an `SR Research Eye Link 1000` (SR Research, Ottawa, Canada) eye tracker. In addition, we recorded the vertical and horizontal electro-oculograms (EOG) by two pairs of electrodes placed above and below, and lateral to both eyes, respectively. The ground electrode for EOG and ECG was attached behind the left ear. Head localization coils were attached to the nasion, and to a set of ear molds that were placed in the left and right ear shell. Participants were provided with neckbands to support their head position, and instructed about ways to minimize movement. After being seated in the scanner they had ten to twenty minutes to find a comfortable position while other system components were initialized. Light was dim and light conditions were consistent for all participants. Potential unknown magnetization or metallic objects were excluded by visually inspecting the MEG signal while participants were following movement instructions. Communication between participant and control room was sustained with an intercom system and scanner room video monitoring.

After the MEG session we acquired structural MRI scans from our participants in order to construct anatomically realistic volume conduction models and detailed cortically constrained source models. Fiducial positions inside the MEG and MRI were assured to be identical using matching sets of ear molds. Vitamin E capsules, attached to identically shaped MRI earmolds, made visual localization in the T1 scans possible.

#### 2.3.4 Stimulus task

Participants were instructed to fixate in the centre of the visual field (fixation point) and to only blink during the phases where this fixation point disappeared from the screen. They were informed about the purely passive nature of the experiment and about the importance and potential loss of attention throughout the sessions, which they were asked to resist or report so they could recover. At the end of each block one single image was presented. To complete the block, the participant had to decide with a button press whether she had seen this image within the previous block or not. Before starting a new block we asked them to adjust their current position relative to their original one using real-time head tracking which is available at our MEG system (Stolk et al., 2013).

### 2.4 Data preprocessing

Data preprocessing and source reconstruction was done using the `FieldTrip` toolbox for EEG-and MEG analysis (Oostenveld et al., 2011) (build: October 27th 2016, GitHub `——short` hash 458858a, http://www.ru.nl/neuroimaging/fieldtrip). Preprocessing steps in this section are described in the order they were applied to the data.

The data was filtered with a DFT filter to remove 50 Hz line noise and its harmonics at 100 Hz and 150 Hz, with a data padding of 10 s. We then extracted trials from the data and corrected the baseline on a trial-by-trial basis by subtracting the average of -250 ms to -50 ms before stimulus onset. A trial was defined as the time period spanning the presentation of the fixation dot, that is from -400 ms to 1600 ms around image onset. We downsampled the data to 300 Hz to numerically stabilize the subsequent Independent Component Analysis (ICA). Before ICA, trial summary statistics were visually inspected and trials with overly high variance and kurtosis were rejected. This served as an initial detection of eye blinks and other muscle movements (Delorme et al., 2007). Muscle artefacts were searched for specifically in a second round of visual inspection, for which the data was high-pass filtered at 100 Hz. In addition to the manual rejection, trials were rejected with an automated function when a z-scaled sample from at least one of the channels exceeded a threshold of five units of standard deviation during stimulus presentation. We went through this procedure in a conservative manner and had to reject 2-10% of the trials for each participant due to various kinds of muscle movements. Subsequently, ICA was applied on the cleaned data to identify the cardiac component and residual physiological artefact components (related to eye blinks and movements). These components were identified by visual inspection, and their corresponding sensor topographies were projected out of the data. After this we inspected the data visually once more to reject trials with potential artefacts and movements, with the same procedure as before running the ICA. The cleaned 300 Hz data was then once more baseline corrected with a window of 250 ms to 50 ms before the trial, and as a final step low-pass filtered at 100 Hz. We did not perform motion correction on the MEG data, relying on participants maintaining their position with the help of the head tracking algorithm.

### 2.5 Source estimation procedure

#### 2.5.1 Preprocessing of the anatomical MRIs

The anatomical MRI images were processed to create individual cortically constrained source models, and volume conduction models for source reconstruction. To this end, we registered the T1-weighted MRI images to the CTF coordinate system by manually locating the fiducial markers. The right hemisphere could be identified with an external additional marker in the scans for all participants but one, for which we reverted to the orientation of the majority of scans.

For the creation of the source models we used FSL, FreeSurfer and HCP workbench. We performed skull stripping on the T1 scans with the Brain Extraction Tool (BET) (Smith, 2002) of FSL, with the threshold set at 0.5. We then ran the FreeSurfer surface reconstruction pipeline up to the segmentation of the white matter. White matter segmentations were visually inspected for artefacts, e.g. slabs of dura being misclassified as white matter. We manually removed such misclassified voxels from the segmented images with a visual tool. Subsequently, the surface-based part of the standard `FreeSurfer recon-all` pipeline was run. These steps resulted in a high resolution left and right hemispheric surface mesh for each participant. These meshes were subsequently surface-registered to a template mesh with 164,000 vertices per hemispheres, and downsampled to a resolution of 4002 vertices per hemisphere, using HCP workbench. We excluded vertices in the midline (corpus callosum and brainstem) for further analysis, so our final meshes contained 7344 dipole locations across both hemispheres.

We created a single shell volume conduction model, describing the inner surface of the skull, using standard `FieldTrip` functionality. The forward model (leadfield) was computed using the method as described in (Nolte, 2003).

#### 2.5.2 Anatomical parcellation labels

In order to get an idea of the anatomical structures corresponding to the sources we matched individual sources on the meshes with anatomical structures from the cortical parcellation from (Glasser et al., 2016) as available in Human Connectome Workbench (Van Essen et al., 2013).

#### 2.5.3 Source activity reconstruction

Single-trial time courses of source activity were computed using Linearly-Constrained Minimum Variance (LCMV) Beamformers (Van Veen et al., 1997) using `FieldTrip`. This solution to the ill-posed inverse problem assumes independent activity of individual sources. It obtains the spatial filter ***s***_*i*_ of a source by solving

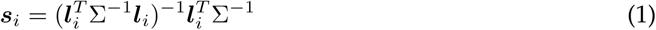

where Σ is the sensor data covariance matrix and ***l***_*i*_) are the elements of the lead field matrix that are estimated for each dipolar source position. The covariance matrix was estimated on all trials and regularized with a diagonal matrix with a value of 5% of the average sensor variance. The location-specific leadfields were norm-normalized in order to account for the beamformer’s characteristic depth bias. The reconstructed 3D dipole moments were projected onto the orientation with the largest variance, leading to one-dimensional time courses for each dipole source.

#### 2.5.4 Source responses

The linear regression models learned to predict the response of a source within a specific time slice after stimulus onset (see Section 2.6). After estimating activity based on the spatial filters and sensor activity X we were left with a sampling rate of 300 Hz per source and trial. To obtain an estimate of the response of each source at a specific time after stimulus onset we binned this signal into consecutive time windows of 30 ms by averaging the signal. These amplitude averages over time windows are subsequently referred to as *source responses* and were used for encoding analysis.

### 2.6 Encoding model

We used the encoding model framework as described in (Naselaris et al., 2011), which tests the similarity between a hypothesized encoding- and brain representation: Input stimuli are transformed into a linearising feature space (the encoding), which is used for predicting source-wise brain activity with linear regression models. Similarity is then estimated by evaluating their predictive power on a held-out test set. As a consequence these feature representations are required to have a direct (linear) relationship with the actual brain activity, which supports interpretability. The linearizing requirement also ensures that there are no implicit nonlinearities learned with more powerful machine learning predictors, i.e. nonlinearities are all explicit within the designed feature space. The power of voxel-wise encoding for both understanding brain representations and for decoding perception from brain activity has been demonstrated before (Naselaris et al., 2011).

#### 2.6.1 Feature transformation

Representation vectors for the stimulus set were obtained in the same way as in (Güçlü and van Gerven, 2015a). As in this paper, we used the pretrained version of the (Chatfield et al., 2014) VGG-S neural network, pretrained on the `imageNet` database (version for `MatConvNet` (Vedaldi and Lenc, 2015)). VGG-S is an *AlexNet-like* network, which means it is similar in architecture to the 8-layer network (Krizhevsky et al., 2012) which was fundamental for the current wave of neural network research. Every stimulus image *n* was passed through the 5 convolutional layers (referred to as layers 1-5) and 3 fully connected layers (referred to as Layers 6-8) of this network. Layer representations of the stimulus image were extracted at the pooling or rectified linear unit (ReLU) layers. The obtained feature maps were flattened to a single representation vector per layer, and then standardized to zero mean and unit variance for each feature across the sample dimension. This hierarchy of representations of an individual image is our hypothesis about the hierarchy of visual system representations during object recognition in the human brain. We assume that this hierarchy can be detected in MEG via the encoding model framework.

#### 2.6.2 Linear model and nested cross validation

For every image n we obtained source responses ***y***_*s,t*_ at a specific time bin *t* and source *s* in each trial in which *n* was presented. We also obtained a feature representation (encoding) ***x***_*n,L*_ of this image *n* in each CNN layer *L*. We estimated independent linear kernel ridge regression models individually for every source, time point and deep network layer using a nested cross-validation routine. Ridge regression was chosen to avoid overfitting these linear regression models. Let ***y***_*s,t*_ be a vector containing responses for every image, for one specific sensor at one specific time bin. Let *X_L_* = (***x***_1,*L*_, …, ***x***_*N,L*_)^*T*^ be a *N* × *M* matrix representing the representation vectors for each image in a specific layer *L*. The linear model’s regression coefficients for a specific time bin, source and layer 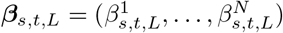 are then estimated by:

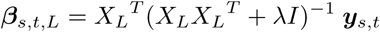

where λ ≥ 0 is the regularization strength in ridge regression. Note that to speed up model estimation we used the kernel formulation of ridge regression.

The performance of each linear model is defined as the Pearson’s correlation between the measured source response and the response predicted by the model. Since we created models for each time point and neural network layer, each of the 7344 sources has 53 × 8 = 424 regression models and thus prediction-activity correlation values.

These models were trained and evaluated with nested 10-fold cross validation. Folds were selected at random from the 950 training trials. Nested within each of these 10 cross-validation runs was one 5-fold cross validation which was used for selecting the regularization strength λ (best λ out of k=5 folds chosen with the procedure described in (Güçlü and van Gerven, 2014)). The prediction-activity correlation coefficients on this best λ fold were used to determine the statistically significant sources (*p* = 0.01 with Benjamini-Hochberg FDR correction over all sources in a given time slice and layer). The final selection of significant sources had to survive significance testing and FDR correction in each of the 10 parent cross-validation folds. Within this procedure, for a single source, performance was estimated by taking the average of the prediction-activity correlations across all folds. These correlation values on the estimation set were used to determine the best-explaining layer for each source, and for selecting those sources that can be explained with higher correlations.

For the other (quantitative) results, encoding and decoding performances were estimated on the validation set, with the selection of significant sources and best-explaining layer per source estimated as described above in the nested cross validation. The models were retrained on all data from the single-trial estimation set for predicting responses on the validation set.

#### 2.6.3 Pixel space control model

Pixel-level models are known to be correlated with visual system responses (Schoenmakers et al., 2013). We used a pixel-level model as a control model. We resized the images to square images of 96 × 96, transformed them to the `CIE 197 6 L*a*b*` color space and took the flattened luminance (L) channel as a representation. In this representation vector spatial coherence between image columns is lost. The information contained in early VGG-S layers will still be similar to this pixel space, but we expected that the larger receptive fields in higher convolutional and the abstraction in fully connected layers outperforms the pixel space model, especially in extrastriate regions. Our results indicated that this control model was not able to predict the responses of a substantial amount of sources for any of our subjects above chance level.

#### 2.6.4 Decoding

We performed a decoding analysis (identification) based on our encoding models. Decodability of an image was defined based on Pearson’s correlations between the predicted and measured source responses to this image in a subselection of sources. The 50 images from the validation set were used for decoding. An image was called *identified* if it obtained the highest prediction-activity correlation, compared to the correlations of all other images. An image was decoded *within the top-5 choice* if its correlation was among the five highest ones.

We retrained encoding models *β_s,t_* for sources and time points on the full estimation set, using information about significance (source-wise best-explaining layers and source-wise average prediction-activity correlations) from the nested cross-validation on the estimation set. We further selected sources by applying a 0.3 threshold on the average correlations from the nested cross-validation. For these selected sources, we predicted the response to a given image from the validation set. Responses of all sources ***s_n,t_*** at time *t* were compared to the measured responses in the validation set (single initial presentation *or* 10-times average) via correlation then.

## 3 Results

This section contains encoding and decoding performances on an individual participant basis and summary statistics on the group level. On the estimation set we estimated the encoding model, the selection of predictable (above-chance) sources, and the optimal layers per source. Encoding and decoding performances are shown on the validation set.

Each encoding model learned to predict the average source amplitude within a specific *τ* = 30 ms time window after image onset. This time bin-specific averaged activity is referred to as *source response.* Models were independently trained in a mass-univariate manner, for each source (whole-brain) and each time point within a trial. This means each of these linear models predicted the activity of a single source (source-wise) in a single time window after stimulus onset (time-wise). The independent variables of these regression models were the 8 different representations from the 8 layers of VGG-S. These present our hypothesized representations for the visual perception of static images. The underlying assumption is that predictability of visual system source responses with regularized univariate models indicates that the tested artificial representations match the biological ones in the visual system.

### 3.1 Encoding of MEG source responses

#### 3.1.1 Encoding performance

Figure 2 shows individual mean encoding performances across all the considered models for predicting source responses. This encoding performance was measured - as everywhere in this manuscript - based on how accurately a model for one source-timepoint combination could predict source responses on the validation set. We could predict brain activity based on the CNN representations for most participants. For the top four participants (4, 12, 1 and 8) we show individual results in the following. Maps for the remaining participants can be found in the supplementary material.

**Figure 2:**
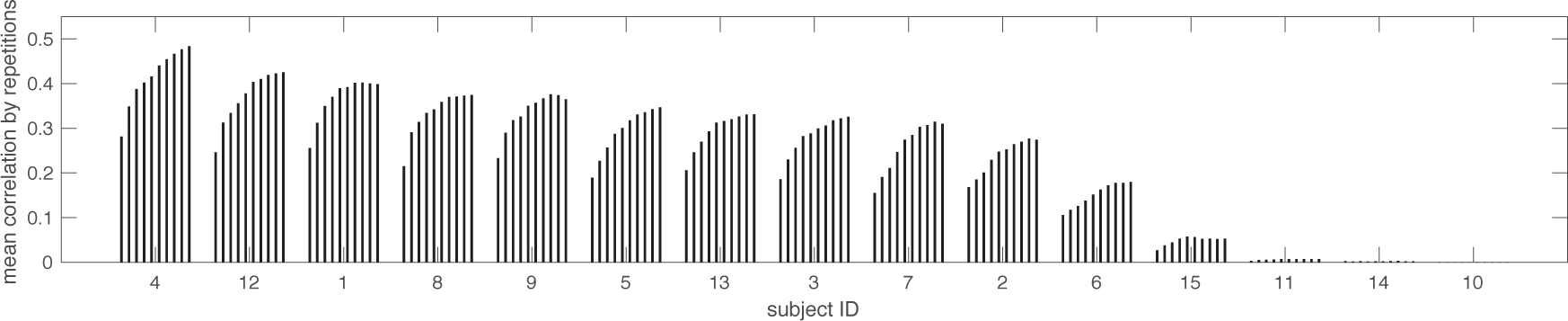
**Mean encoding performance** in relation to the number of repetitions on the validation set for sources anatomically assigned to the visual cortex. The model shows considerable mean correlations for 10 out of 15 participants. For 3 participants it is not predictive at all, and for 2 average correlations are low. We predicted the activity for each source for the time bins between 75 ms and 600 ms after image onset for the 50 images in the validation set (before the 75-105 ms time bin, for most subjects no source activity can be predicted). Predictive models were trained for each source-time bin combination on the full estimation set; with significance and optimal layers estimated during cross-validation within the estimation set. Correlations between the predicted and the measured responses per source and time bin were then taken across validation set stimuli. The mean shown here summarizes these correlations for sources assigned to the visual system areas with our anatomical parcellation. The increased SNR from averaging over repetitions improves encoding performance. However for most participants the average over 10 repetitions appears to be close to a performance plateau.

The magnitude of the prediction-activity correlations on the validation set of the models explaining a source best is shown in Figure 3. As in (Güçlü and van Gerven, 2015a), the majority of high correlations was observed around the early visual cortex. From a temporal perspective we observed the highest correlations during the first 75-135 ms after image onset, and the magnitude diminished over time. Some participants already showed predictable activity as early as 45-75 ms after image onset.

**Figure 3:**
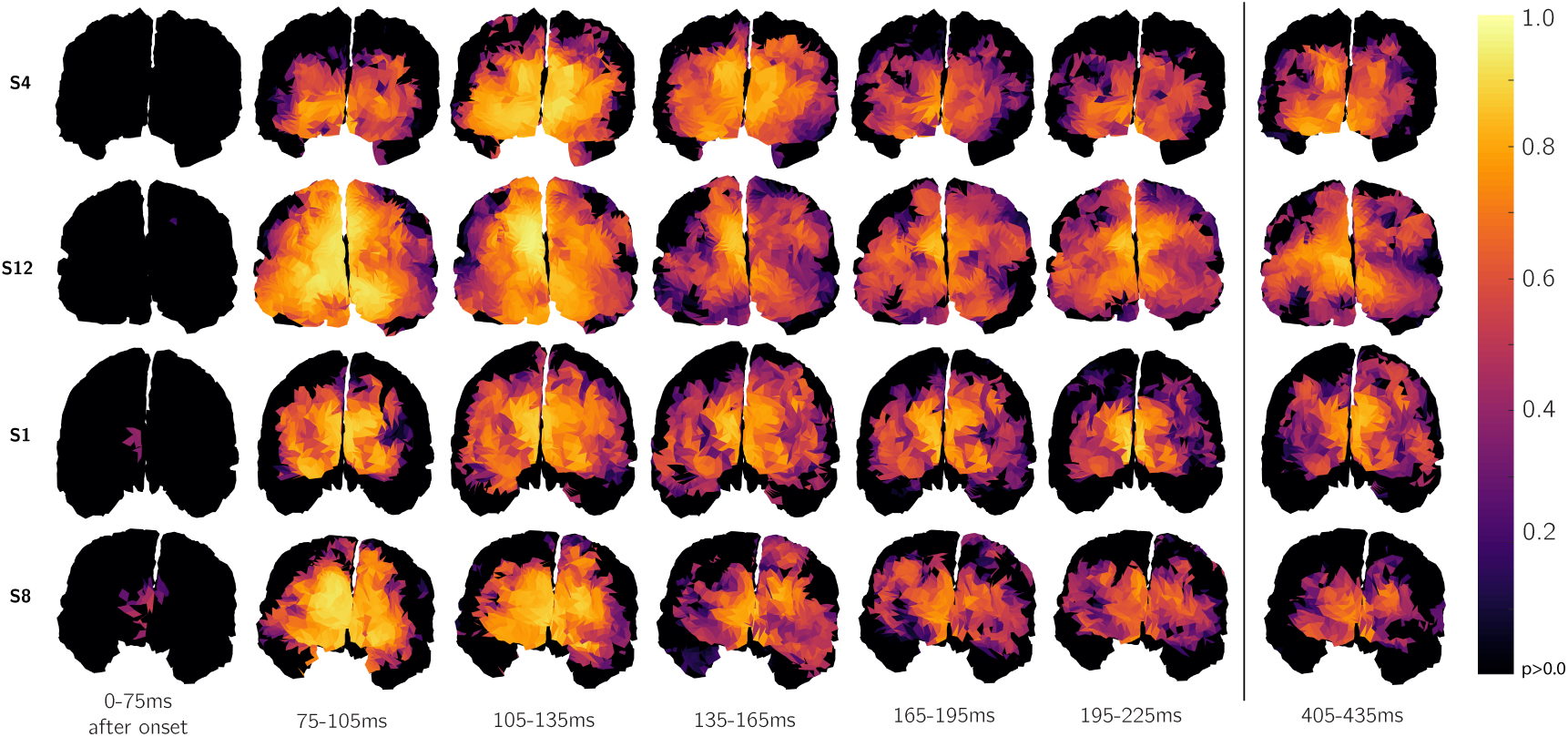
**Prediction-activity correlations over time** on the 10-times validation set for the encoding models using the best-explaining layer as in Figure 4. Views are centered at the occipital lobe. We show all above-chance correlations. We observe the highest correlations around the early visual cortex, and lower correlations in extrastriate areas.

#### 3.1.2 Representational gradients

Figure 4 shows which of the VGG-S layers can explain each individual source best for our top participants. Significant sources and best layers for source-time point correlations were determined within nested cross-validation on the estimation set. In this sense, Figure 4 is similar to Figure 2A from (Güçlü and van Gerven, 2015a), adding views on the temporal dimension. After training the model on all sources across the cortical surface, we observed for all participants that only source responses across and around the occipital lobe could be predicted above chance level (*p* < 0.01, Benjamini-Hochberg FDR). In addition to this, Figure 4 only shows sources where the best layer resulted in a prediction-activity correlation of more than 0.3 on the estimation set, observed as an average of the correlations from the top-level cross validation folds. This threshold on the mean correlation over folds was used for a subselection of results in order to focus on sources that reach higher effect sizes, which can safely be called a meaningful similarity of representations, instead of merely passing the significance threshold.

**Figure 4:**
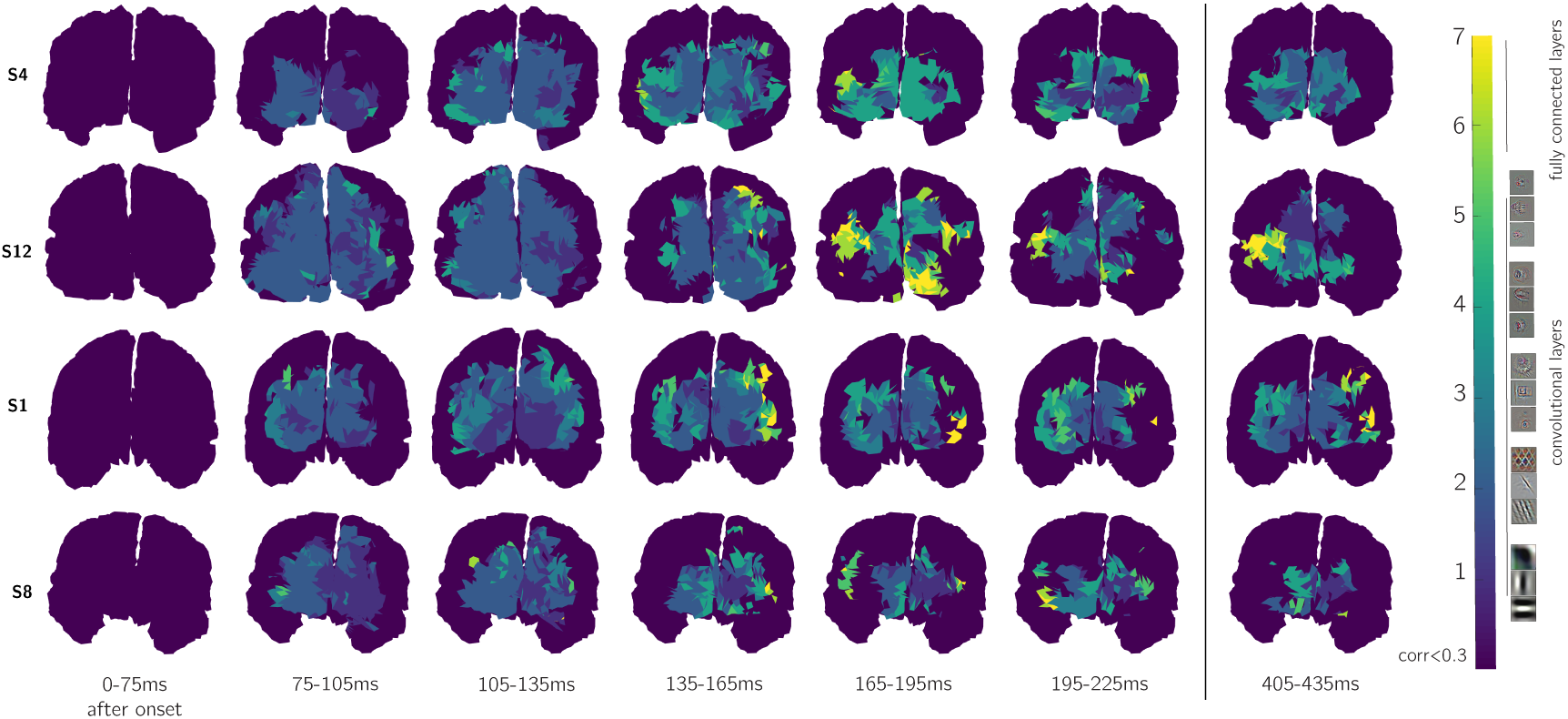
Source-wise layer assignments over time for four participants. Views are coronal from the posterior onto the occipital lobe. Each source-time bin combination gets assigned the representation layer that explained it best during the nested cross-validation on the estimation set, measured in the average correlation across all folds. Sources are shown on individual brains based on the Freesurfer models. Non-significant sources and significant ones with low correlations (<0.3) are discarded in these maps. The manifestation of the fully-connected layers 6 and 7 first occurs after 135ms for most participants. Before this time convolutional layers are expressed, starting with a widespread manifestation of layer 1 and 2 in the early visual cortex region. After the expression of fully connected layers for some, but not all participants we see sustained activity, here shown for the time bin 405-435ms. The colormap was chosen to reflect the division between convolutional and fully-connected layers.

The maps show that all layers are expressed before the time slice starting at 195 ms for adjacent sources in one or more specific regions. In time, the cascade starts with Layers 1 and 2, which cover a widespread region of the occipital lobe within 75-105 ms. Layer 2 can explain more sources than Layer 1 for all participants at this early stage. The features of this second layer are texture-like combinations of Gabor features, for instance lattices, and have slightly larger receptive fields than Layer 1. Layer 1 in contrast predominantly represents small Gabor-like features, resembling striate cortex receptive fields. The nature of the MEG signal and how we recorded and analyzed it may not lead to detecting single striate cortex receptive fields, which could be the reason that the compositional nature of Layer 2 is a better explanation for striate cortical surface responses.

The layers progress towards extrastriate regions that can be predicted more accurately with higher convolutional (3-5) layers. The variability of the maps across participants is striking. It is noticeable that several sources that could be explained best with the low convolutional layers in previous time slices are best explained by higher layers in later time steps.

In the time slices between 135-195 ms, Layer 6 and 7 are first expressed. These are the first fully-connected layers, which means that they lose most spatial information after the highest convolutional layers. They are the most abstract representation of an image in VGG-S.

Note that as in (Güçlü and van Gerven, 2015a) the representation from the softmax layer 8 is unable to explain any source optimally. Only few sources pass the significance threshold when predicting with it, resulting in very low prediction-activity correlations. This is not surprising, since the layer represents a probability distribution over the 1000 categories of the ImageNet competition, on which VGG-S was originally trained. ImageNet categories are not defined in order to reflect real-world categories exhaustively. E.g., a large subset of the 1000 categories represents a specificity test on fungus and dog species. It is possible to change the layer 8 semantic categorical representation by fine-tuning the network towards a different ground truth of categories, however for comparability we chose to stay with the original state of the network.

Figure 5 shows the mean (A) and median (B) onset times for each VGG-S layer. It is equivalent to showing the layer onsets visible in Figure 4 in time. Note that due to our encoding procedure, the results presented here have no higher temporal accuracy than 30 ms (*τ*), so onset times were centred between the time boundaries and Sheppard’s correction was applied to the standard deviations.

**Figure 5:**
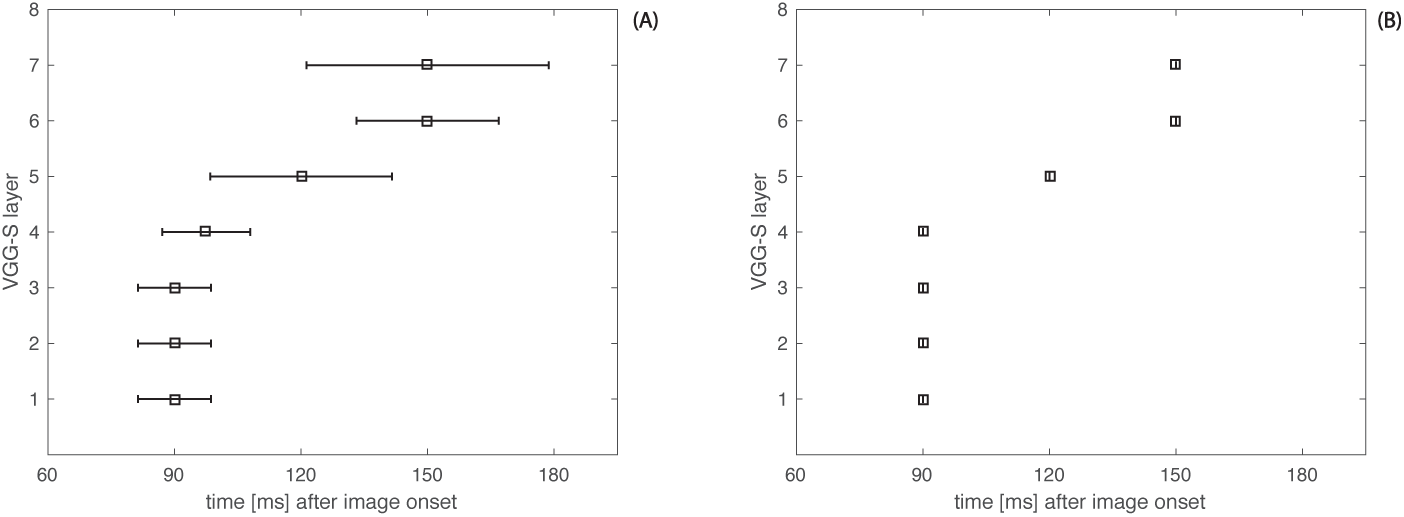
**Temporal onset of layers**, average (A) and median (B) over the 12 participants for which the encoding model was predictive. Shown is the first time bin in which the respective layer explains at least one source best with correlations above 0.3. We indeed observe that the hierarchy from Layer 1 to 7 is being expressed sequentially within 200 ms after stimulus onset. The first four convolutional layers first occur within the 75-105 ms time slice.

We observe that the layer cascade follows the hierarchy from VGG-S. Across participants all layers are traversed within 200 ms after stimulus onset, where the mean time needed to express the most abstract Layer 7 is 150 ms. There is a temporal gap before and after Layer 5. Note that the convolutional feature detectors in Layer 5 are special in that they already express complex templates of full objects. Incorporating the special role of Layer 5, there is a temporal division between lower and mid convolutional layers (Layers 1-4) and the most abstract layers (Layers 5-7). This is most apparent in the median (B).

MEG has higher spatial resolution than other direct non-invasive measures of neural activity due to less skull distortion, which allows much more accurate localization of the sources of measured activity. While the spatial resolution is nevertheless limited, the higher resolution can lead to more accurate localization of the anatomical origin of source activity. We linked the source meshes to the recently introduced anatomical parcellation by (Glasser et al., 2016) using the `Human Connectome Workbench software` (Van Essen et al., 2013). The 181 anatomical labels for each hemisphere were clustered into the 22 top-level regions defined in the supplementary material of (Glasser et al., 2016), of which we focus on regions around the occipital lobe active in visual processing (see Table 1).

**Table 1:** **Clusters of top-level anatomical regions**, according to the supplementary material of (Glasser et al., 2016). Each source was assigned the label of its corresponding template region.

| Top-level region | Anatomical labels from (Glasser et al., 2016) |
| --- | --- |
| Primary visual cortex | V1 |
| Early visual cortex | V2, V3, V4 |
| Dorsal Stream | V3A, V3B, V6, V6A, V7, IPS1 |
| Ventral Stream | V8, VVC, PIT, FFC, VMV1, VMV2, VMV3 |
| MT+ complex and nearby | V3CD, LO1, LO2, LO3, V4t, FST, MT, MST, PH |
| Inferior parietal cortex | PGp, PGs, PGi, PFm, PF, PFt, PFop, IP0, IP1, IP2 |
| Posterior cingulate cortex | DVT, ProS, POS1, POS2, RSC, v23ab, d23ab, 31pv, 31pd, 31a, 23d, 23c, PCV, 7m |

Using this strategy, we could gain insight into the anatomical regions where deep neural network layers were expressed over time. In Figure 5 we observe that the lower convolutional layers (1-4) dominate across all regions early in time. In contrast, the most abstract layers (6, 7) are expressed in and around the ventral stream regions after 135 ms post-stimulus onset.

#### 3.1.3 Specificity of low- and high-level layers

We investigated whether the different nature of convolutional and fully connected layers affected the predictive power in space and time. For this we contrasted the spatial distributions of explanatory power of Layer 2 (a low-level convolutional layer, covering more sources than Layer 1), Layer 4 (a high-level convolutional layer that has not reached the object template representation as expressed by Layer 5), and Layer 6 (the first fully connected layer). Visual and quantitative results can be found in Figure 7 and Figure 8. Layer 4 (and other higher convolutional layers) spread towards extrastriate regions, while showing similar predictive power as Layer 2. The sources that could be explained by the fully-connected Layer 6 are, in contrast, more separated from those that can be explained by Layer 2. Figure 8 shows how this division evolves over time. The reason for this divide could involve different representational properties of the contrasted representations: The fully-connected layers lose most spatial information and can therefore be considered more translation-invariant. They also carry the most abstract representations needed for object identification. Convolutional layers have localized responses, and carry mostly low-level structural information. The contrasts presented here provide another indication that higher layers (both convolutional and fully-connected) indeed explain extrastriate visual system activity.

**Figure 6:**
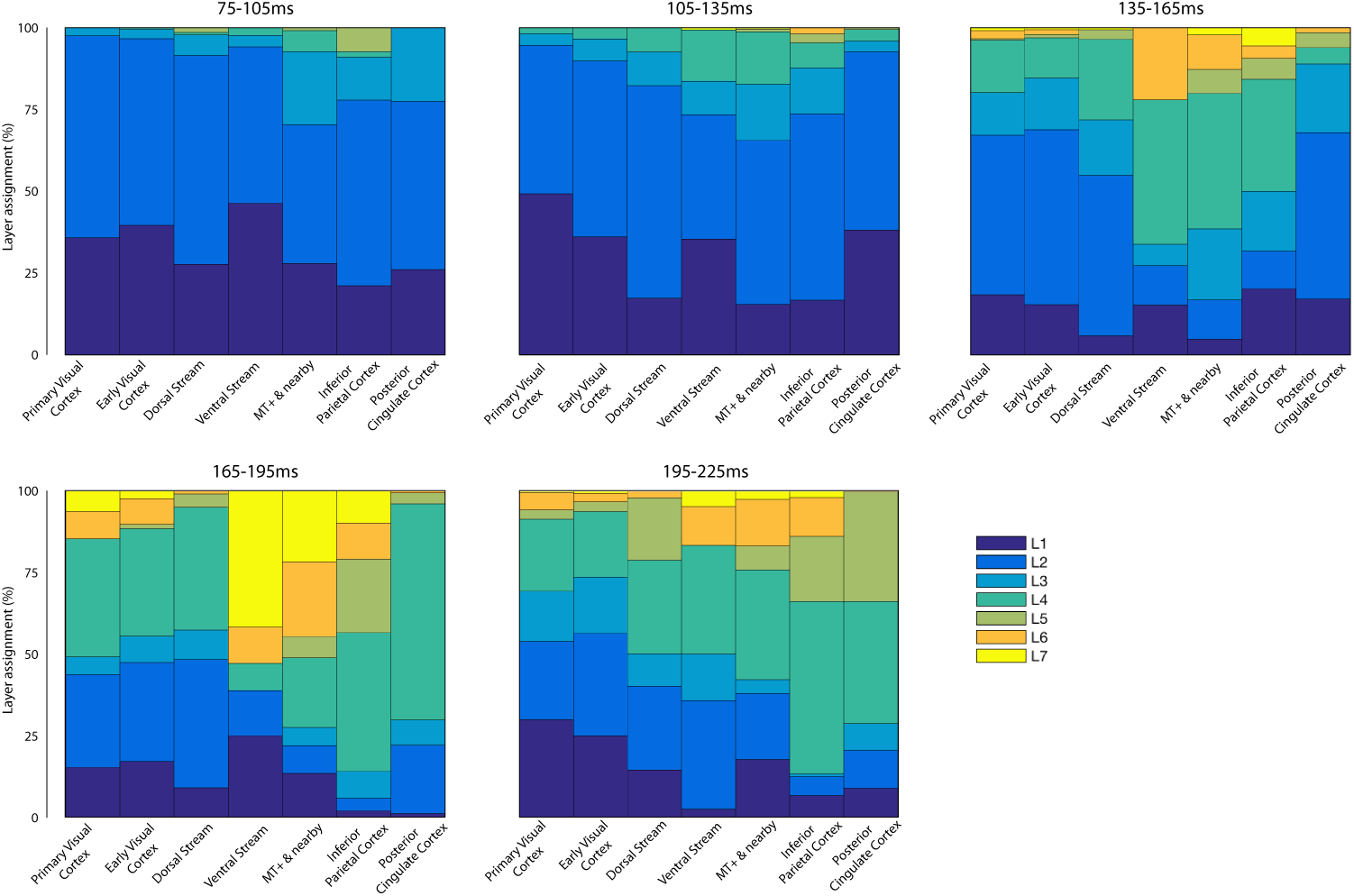
**Number of sources in an anatomical region assigned to the network layers**, averaged over participants 4,12,1, 8. Early convolutional layers up to Layer 4 are expressed across all layers. The most abstract Layers 6 and 7 appear in the ventral stream and neighbouring regions after 135 ms (sources with above 0.3 correlations).

**Figure 7:**
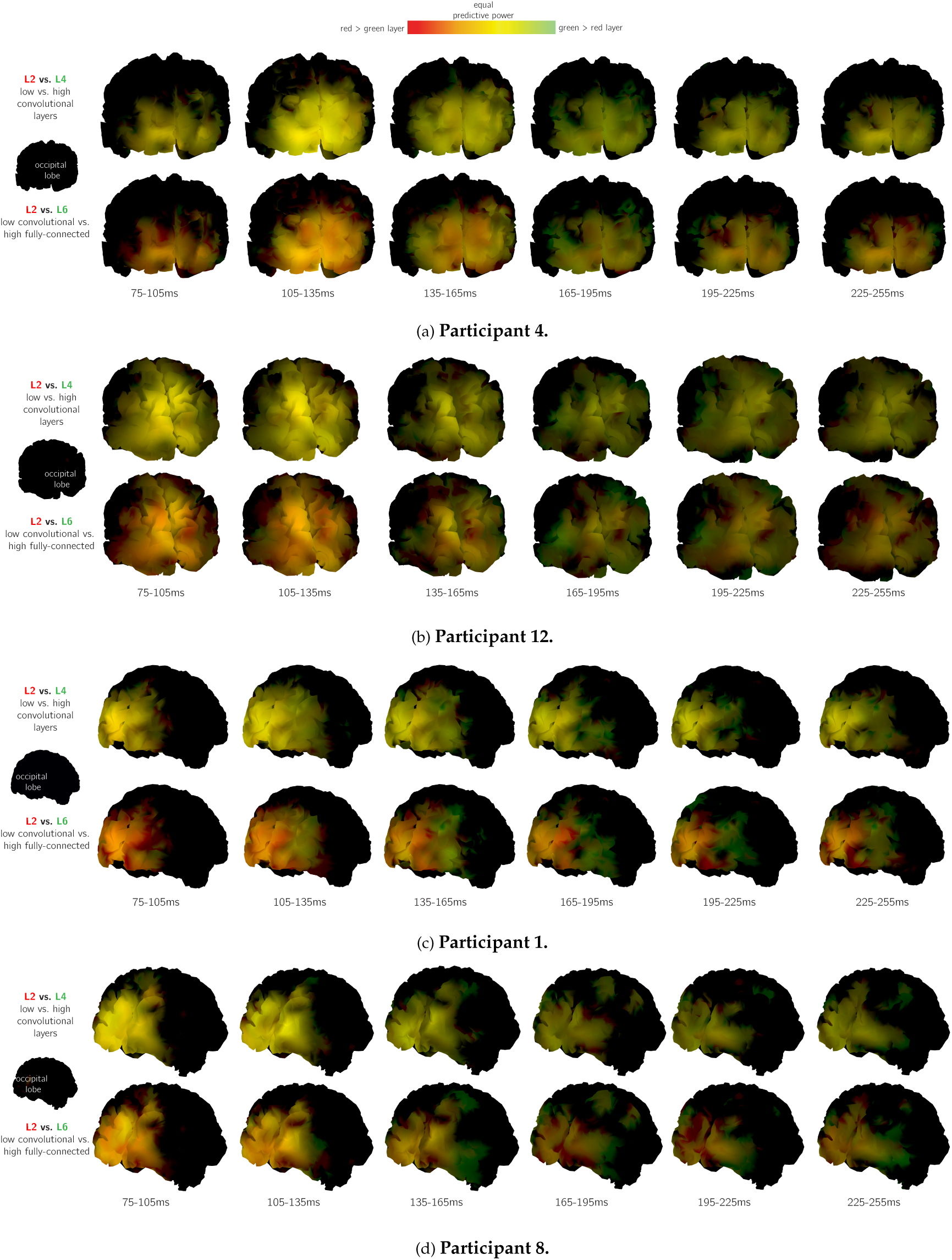
**Spatial distribution of predictive power of convolutional and fully connected layers** over the first 255 ms. Sources explained by early convolutional layers and fully connected layers do not appear in the same regions. Convolutional layers explain similar regions, with the mid-level convolutional layers spreading out into extrastriate areas. For the visualization, correlations for each layer are normalized by the highest correlation observed for each participant and Fisher-z corrected to allow linear comparability. Correlation values for the given layers then fill either the red or the green RGB color channel, highlighting sources were one layer outperforms the other, and leading to a mixture (yellow) if both layers can explain a source equally well.

**Figure 8:**
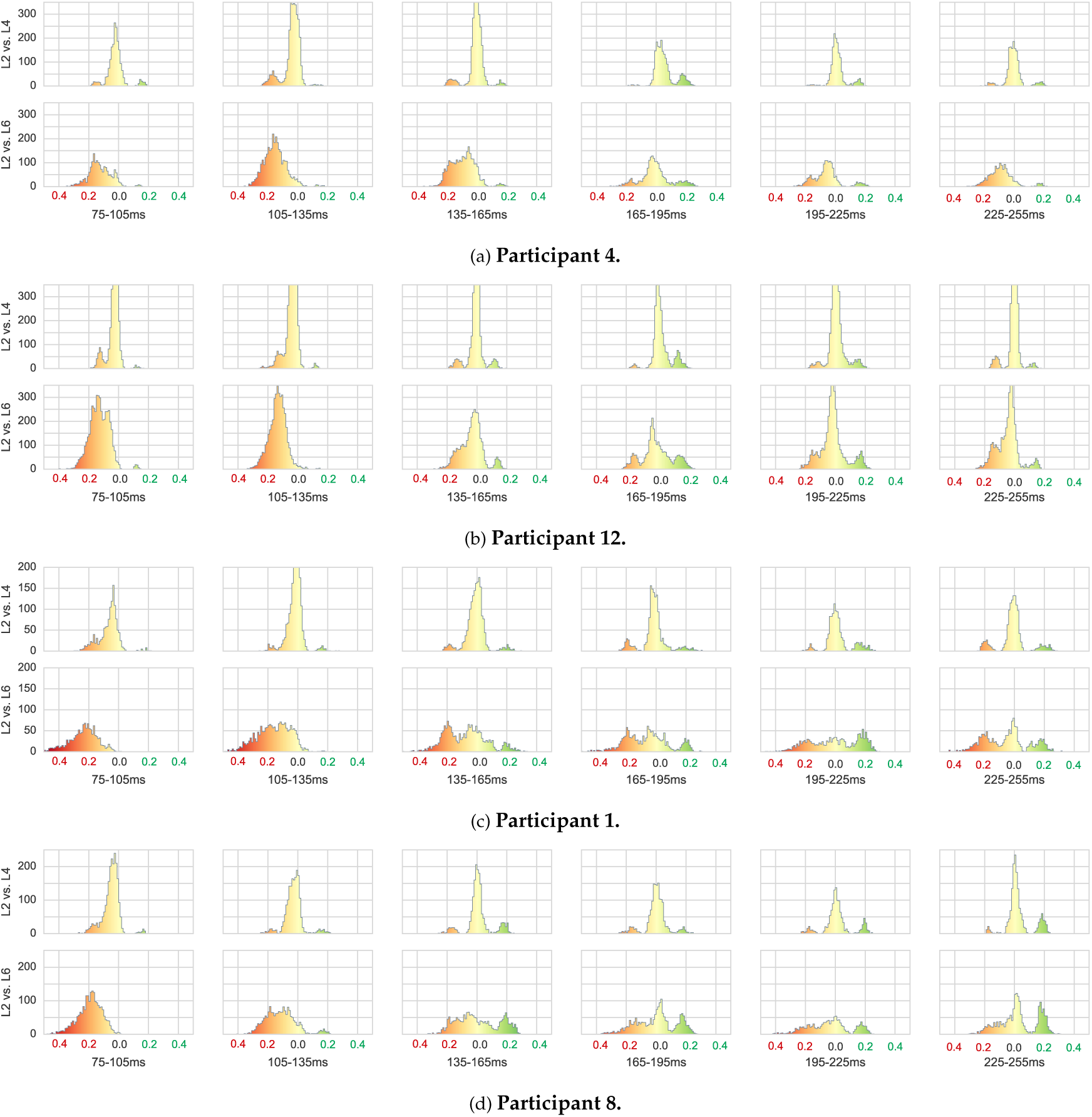
**Histograms of source-wise correlation differences between higher and lower layers** 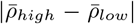. Fisher-z corrected correlations were used to allow linear comparability, and normalized by the highest observed correlation for each participant (close to 1.0 for all). Sources with low prediction-activity correlation or with similar predictability appear around 0 while source better explainable by either red or green layers in the left or right modalities of the histogram respectively. We observe the progressing division over time into low and high layers that we also saw in Figure 7. The division is more pronounced when comparing the low convolutional to fully connected layers. Comparing only convolutional layers leads to more sources in the central modality that represents similar explainability. This indicates a spatial division into regions with low-level information (including spatial) and high-level, abstract information (fully-connected, translation-invariant).

### 3.2 Decoding from MEG source responses

The 50 images from the validation set were used for decoding. All encoding models were retrained on the full estimation set using significant sources and optimal layers determined by the nested cross-validation procedure. Source responses were predicted for the 50 images in the validation set and pairwise correlations with all measured source responses were created. Figure 9 presents these pairwise correlations^1^ for participants 4, 1, 12 and 8, using averages over ten repeated validation set responses within 75-225 ms after image onset. The time slice corresponds to the time that is needed for traversing the neural network hierarchy. We again selected the sources that reached higher correlations than 0.3 within the nested cross-validation on the estimation set. The anti-diagonal corresponds to the correlations between our predictions and the observed activity for these stimuli. The magnitude of these diagonal correlations indicates whether images from the validation set can indeed be identified using predictions from the encoding models.

**Figure 9:**
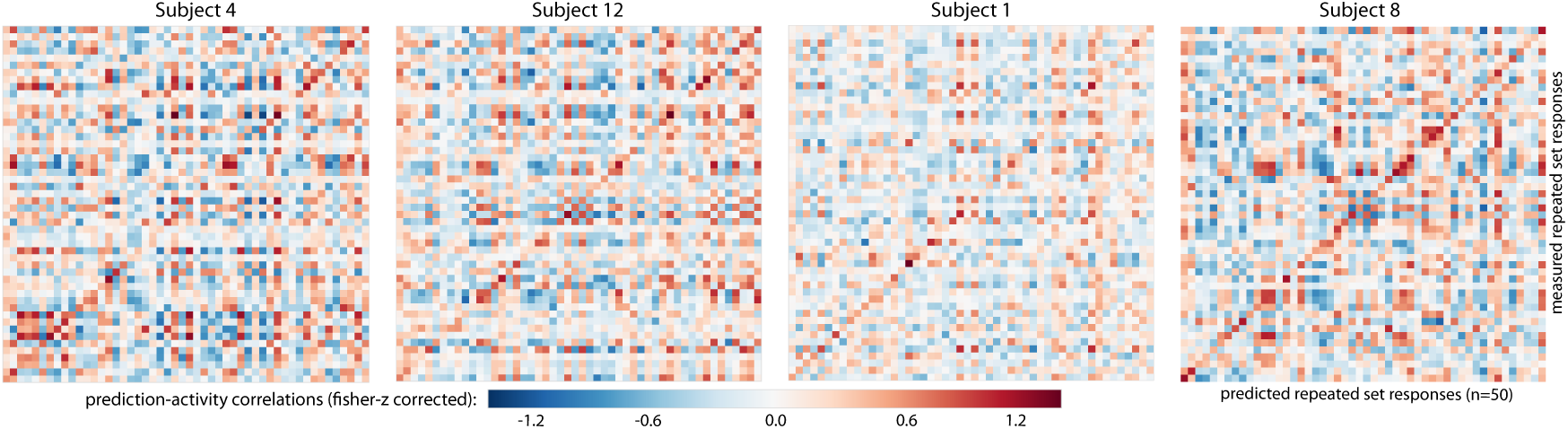
**Correlations (on resampled data) between predicted and measured averaged activity** for four participants for the 50 images of the validation set, between 75 ms and 225 ms. Sources that reached higher average correlations than 0.3 during nested cross-validation on the estimation set were selected for decoding. Predicted source responses were compared to measured responses resampled over ten repetitions.

Similarly, Figure 10 shows these correlations for the same participants for single-trial responses, again for the 50 images in the validation set. As expected the correlations between the predicted and observed responses to a target image are lower, but still indicate decodability.

**Figure 10:**
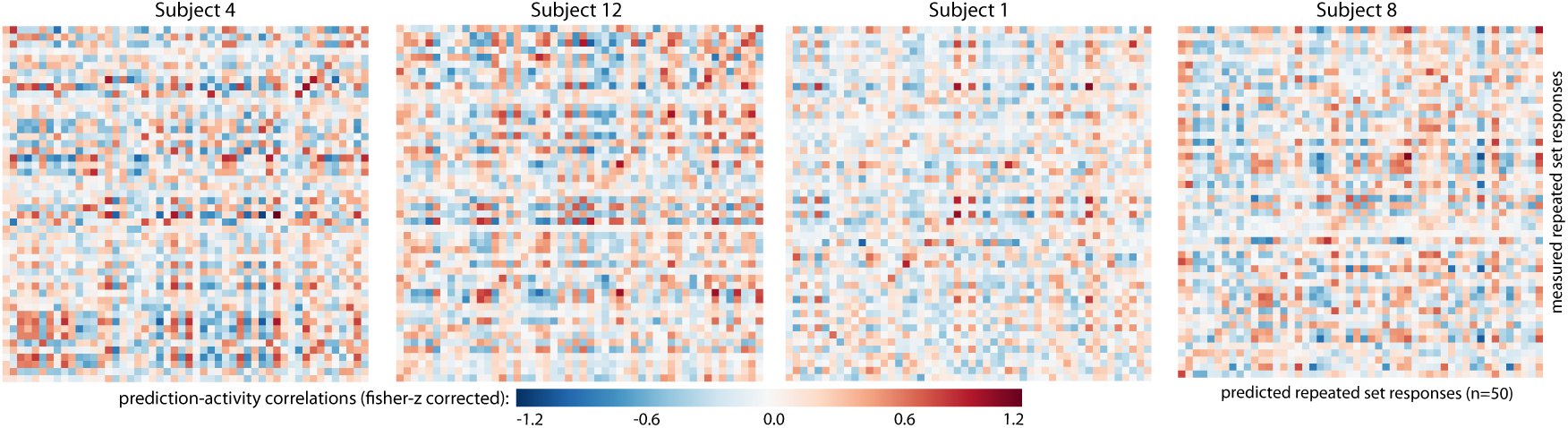
**Correlations (single trial) between predicted and measured averaged activity** for four participants for the 50 images of single first trials from the validation set, between 75 ms and 225 ms.

Figure 11 shows the number of images that can be either *identified* or for which the presented image belongs to the *top-5* most correlated images, for every participant. For most participants, this number of *identified* or *in top-5 choice* images is above chance level (2% and 10% respectively) at most time points during image presentation. Up to 70% of the presented images can be found within these top-5 choices, depending on the participant. For a few participants this procedure fails however, and we observe no trace of the learned model in the data. We know that some of our subjects were not able to maintain their head position throughout a block, and it is likely that not all of them were capable of focusing on passively viewing objects for a long time.

**Figure 11:**
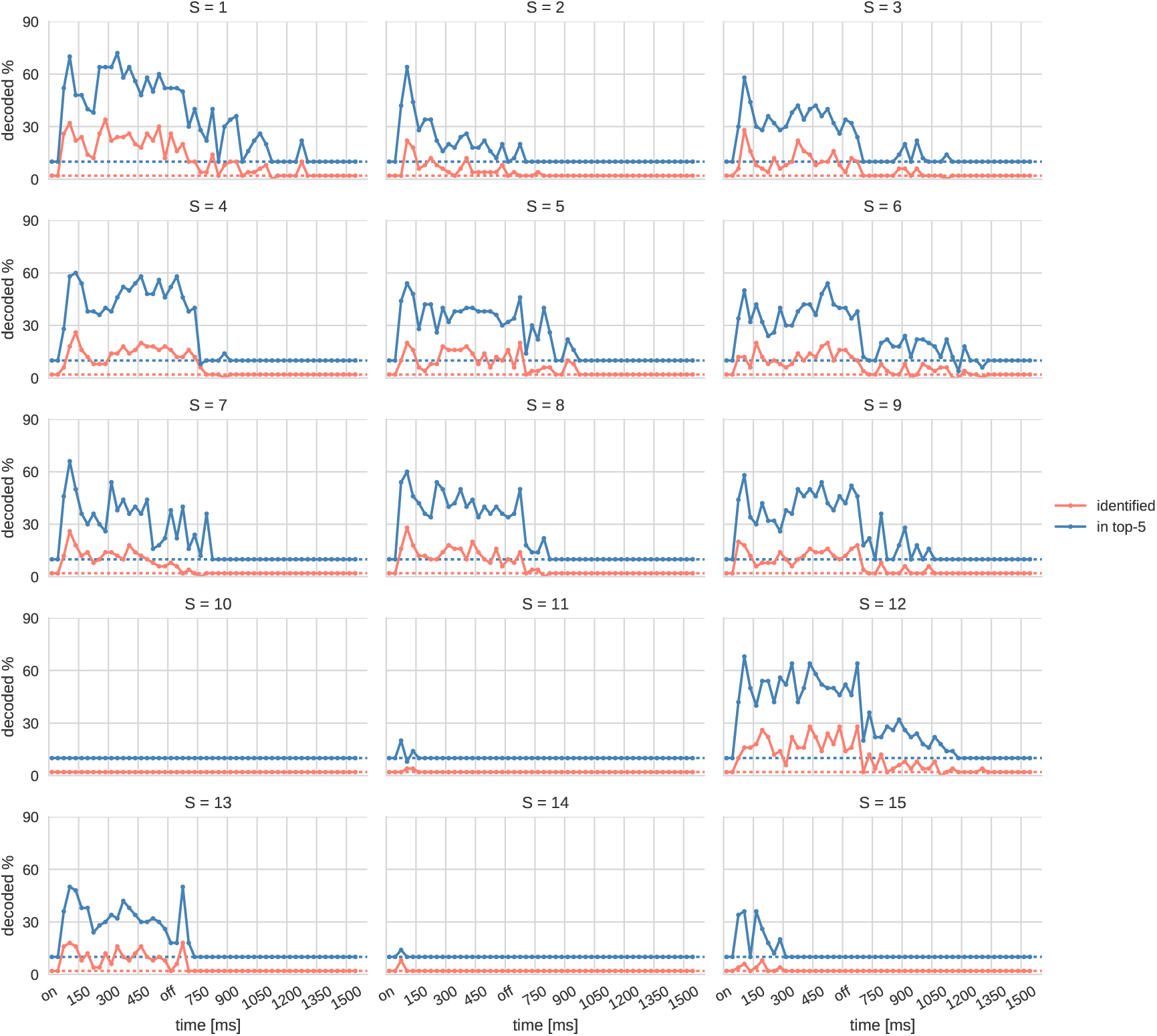
Decoding performance over time for repeated trials. Predictions of source responses given a stimulus image using the significant sources and best layer per model as estimated on the estimation set are correlated to actually measured activity (averaged over the 10 repetitions). The red line shows how many images could be directly identified; that is the correct image achieved the highest correlation. The blue line shows how often the correct image was among the top-5 correlated images (*top-5 choice*). The dotted lines represent the chance levels (identified: 2%, within top-5 choice: 10% for the 50 test set images).

The average of the decoding performances from Figure 11 over the 11 participants showing above-chance decoding performance can be found in Figure 12. On average up to 20% of the presented images can be identified in these participants, and in up to 58% of the cases the images belong to the top-5 most correlated images. For single-trials these numbers decline to 42% and 18% respectively (see Figure 13). After an initial peak of decoding performance, it declines during the initial 225 ms. After 300 ms, while the image is still on the screen, there is an increase in decoding performance again.

**Figure 12:**
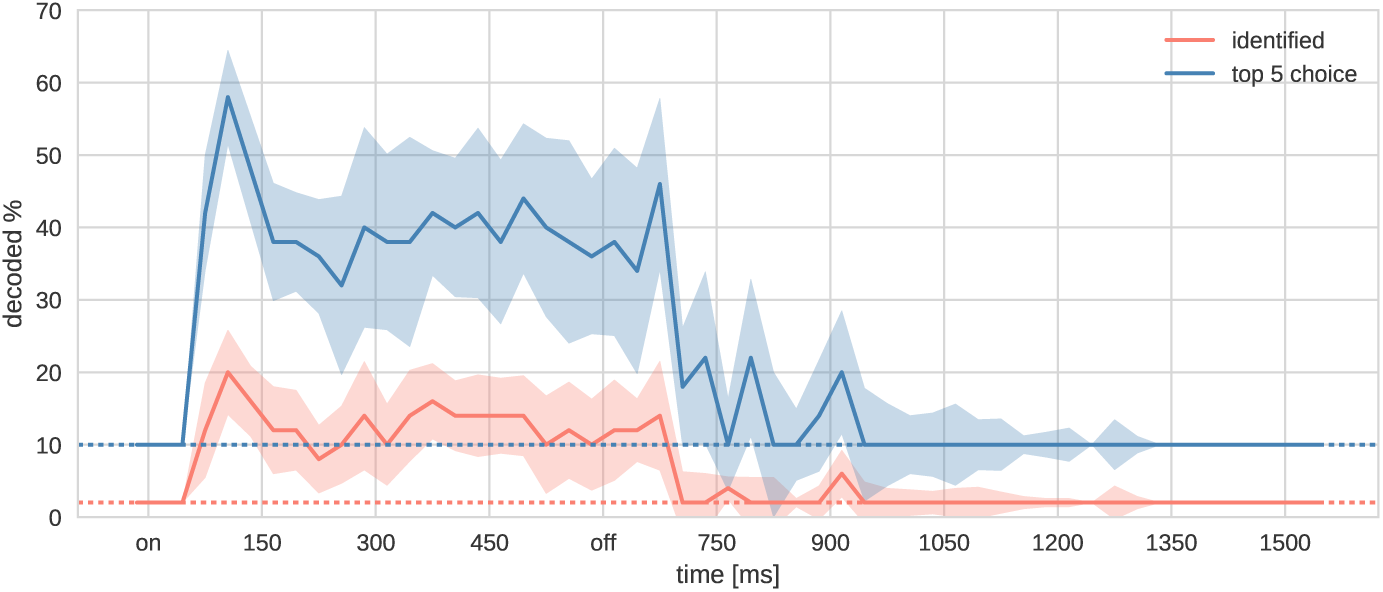
**Mean decoding performance over time** for participants that showed above-chance decoding performance in Figure 11. Error bars present the standard error of the mean across participants. Dotted lines represent the chance levels.

**Figure 13:**
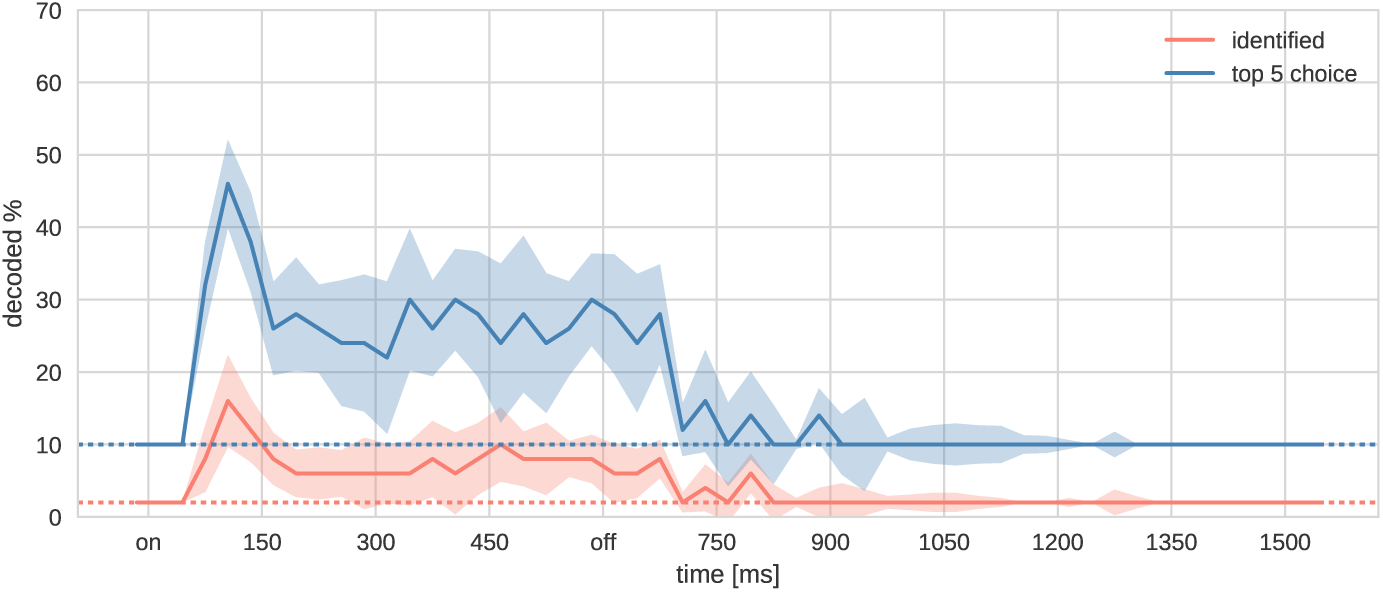
**Single-trial mean decoding performance over time** for participants that showed above-chance decoding performance in Figure 11. Initial presentations of the 50 images in the validation set were taken, so this is not influenced by diminished attention through repetitions or late blocks. Error bars present the standard error of the mean and dotted lines the chance levels.

All participants whose brain activity can be decoded above chance level show sustained de-codability over the whole image presentation period. Participants 1, 6, 7, 9 and 12 also show above-chance decodability after stimulus offset. For them the top-5 choice decoding rate is still around 30% long after the image disappears at 600 ms. Some of these participants also show up to 10% above-chance identification rates in this delay window.

## 4 Discussion

In this study, we show that stimulus representations from a deep convolutional neural network can predict MEG source activity across the visual system, both in space and time. Earlier layers of the network predict neural activity as early as 45-75 ms after stimulus onset in early visual regions, whereas higher layers predict later activity in higher visual regions. Encoding performance as a function of the number of repetitions indicate that 10 repetitions are close to a performance plateau for most participants (cf. Figure 2). The observed feed-forward activity sweep is completed around 150 ms on average, closely matching other studies on the temporal properties of object recognition (Thorpe et al., 1996). In addition to this, the temporal sequence of representations is indeed close to the hierarchy which AlexNet-like networks such as VGG-S learn. Furthermore, the anatomical association of the translation-invariant fully-connected abstract layers with the ventral stream matches earlier studies on the localization of object recognition (DiCarlo et al., 2012). Finally, we were able to invert the encoding models for decoding, resulting in far above chance decodability. All analyses could be done in source space with accurate individual source meshes.

Our results are in agreement with earlier results obtained by comparing CNN layer representations with MEG-based neural representations (Cichy et al., 2016; Cichy and Teng, 2016). Notable differences between our approach and previous work are that our results have been obtained by an encoding approach instead of an RSA approach. The encoding approach affords the prediction of individual responses in MEG source space rather than the assessment of representational similarity across time using sensor-level data. Furthermore, the encoding approach affords decoding of neural responses using an identification strategy. Estimation of an encoding model does require an order of magnitude more data compared to the use of an RSA approach.

We tested whether our results were driven by low-level image properties using the luminance channel from the `L*a*b*` color space as a control model. However this did not yield comparable predictive power - depending on the participant, at most only a few sources around the early visual system passed significance thresholds. We could not construct meaningful maps or decode from these alternative encoding models.

An intriguing finding of the present work is that early in time, most regions are predicted well by low-level layers, whereas later in time, most regions are predicted well by high-level layers (Figure 6). Even early visual areas are described well by high-level layers related to abstract semantic features as time progresses. This might point towards a recurrent integration process where semantic or broader structural scene information is distributed across brain regions. At the same time, given the limited spatial resolution of MEG, we cannot rule out that these results are due to a spillover of information encoded in nearby brain regions.

The decoding results indicate that source-space encoding on MEG data can be suitable for more advanced decoding techniques such as reconstruction rather than identification of visual stimuli. Especially when using video stimuli, MEG would not suffer from the low-pass filtering effect of the hemodynamic response function that affects fMRI movie decoding. However, as expected given the nature of the noisier, less localizable MEG signal we do not reach the accuracies demonstrated in fMRI (Kay et al., 2008; Güçlü and van Gerven, 2015a). Also, the above-chance decodability observed post-stimulus offset in some participants indicates that, using a suitable experimental design, we may be able to decode complex stimuli from working memory and imagery using our approach.

As demonstrated by our findings, convolutional neural network models of the visual system yield insight into the spatiotemporal dynamics of neural information processing. Neural networks have been shown to strongly benefit from biologically inspired mechanisms such as local convolutional operations (LeCun and Bengio (1995)), DropOut (Hinton et al., 2012) or ReLU nonlinearities (He et al., 2015). At the same time, there are ample opportunities to improve model fit by taking other biologically plausible principles of neural information processing into account. Future work could focus on neural networks that make use of alternative architectures (e.g., residual and recurrent neural networks (He et al., 2015; Hochreiter and Schmidhuber, 1997)), solve different kinds of problems (e.g. semantic segmentation (Güçlü et al., 2017)), are trained on other kinds of data (e.g. multimodal data (GUclUtUrk et al., 2016)), use alternative objective functions (e.g., maximizing future frame prediction (Mathieu et al., 2015)) and/or use a different learning paradigm altogether (Song et al., 2016; Bosch et al., 2016).

Concluding, we expect new advances that bridge the gap between artificial and biological brains to ultimately provide new insight into the computational basis of neural information processing.

## 5 Acknowledgements

This research was supported by VIDI grant number 639.072.513 of The Netherlands Organization for Scientific Research (NWO). J.-M. Schoffelen was supported by VIDI grant number 864.14.011. We thank our colleagues Marieke van de Nieuwenhuijzen and Luca Ambrogioni who provided comments, insight and expertise that greatly assisted this research.

## Footnotes

1 Due to the nonlinear nature of (especially higher) Pearson’s correlations we show Fisher-z-transformed correlations, transforming all correlations into the range [−∞, ∞] instead of [0, 1]. A Fisher-z corrected correlation of 1.2 corresponds to an uncorrected one of 0.834 and comparisons of difference magnitudes can be made.

